# Cerebellar Contribution to Absence Epilepsy

**DOI:** 10.1101/2020.11.10.376004

**Authors:** Leonid S. Godlevsky, Oleh R. Pinyazhko, Olesya B. Poshyvak

## Abstract

The new aggregate data analyses revealed the earlier missing role played by the cerebellum long-term electrical stimulation in the absence epilepsy. Neurophysiologic data gained by authors favor that cerebellar serial deep brain stimulation (DBS) (100 Hz) causes the transformation of penicillin-induced cortical focal discharges into prolonged 3,5-3,75 sec oscillations resembling spike-wave discharges (SWD) in cats. Such SWDs were not organized in the form of bursts and persisted continuously after stimulation. Therefore the appearance of prolonged periods of SWD is regarded as a tonic cerebellar influence upon pacemaker of SWD and might be caused by the long-lasting DBS-induced increase of GABA-ergic extrasynaptic inhibition in forebrain networks. At the same time, cerebellar DBS high-frequency (100 Hz) suppressed bursts of SWD observed during the phase of stimulation. Different types of cerebellar DBS upon epileptic activity emphasized the absence seizure facilitation discussed with the reviewed data on optogenetic stimulation, neuronal activity of cerebellar structures, and functional magnetic resonance imaging data.

## 1. INTRODUCTION

Childhood absence epilepsy (CAE) represents a benign form of generalized epilepsy, which comprises 10-15% of all childhood epilepsies [1, 2, 3]. Both experimental and clinical absence seizures (AS) represent minimal visible clinical symptoms and typical and profound EEG deterioration [4–8].

Main pathogenetic peculiarities of spike-wave discharges (SWD) as a biomarker of AS confined to hyperexcitation within the cortico-thalamo-cortical (CTC) network [6–9]. Intensively developed investigations of genetic forms of absence seizures performed during the last decades [10–14]. Calcium channel deteriorations may underlay such hyperactive state of neurons within the CTC networks due to an α(1A) voltage-sensitive calcium channel gene mutation – as established in tottering (tg) and tg(la) mice [14, 15, 16, 17].

Recent data proved that a pathological enhancement of GABAergic signaling within a thalamocortical network is necessary and sufficient for nonconvulsive AS development [18–25]. The overwhelming inhibitory GABA-ergic effect is in charge of absence SWD promotion [18, 19, 26]. It might be assumed that increased activation of extrasynaptic GABAA receptors and augmented tonic GABAA inhibition in thalamocortical neurons are in charge of such effect [20].

Continuously developed absence seizures in rats induced via gamma-butyrolactone (GHB) (200 mg/kg, i.p.) administration is characterized by a lower frequency of discharges (2,64-3,59 Hz), which are not organized as bursts [24, 27]. The enhanced GABAergic inhibition serves as a basic conception of AS manifestations, and compounds which intensified it recognized as proabsence ones [25]. The reticular thalamic nucleus (RT) ‘s leading role in the GABA-ergic suppression of specific thalamic nuclei was strengthened last time [28]. Such data were in line with the two-phase effects of vigabatrin upon SWD [29] and with the suppression of SWD caused by DBS of RT [30].

Despite relatively good pharmacological control of CAE manifestations, resistance to treatment is actual [31–34]. Searching for alternative therapy of not-responsive AS targets for DBS proved promising [35]. Among others, thalamic structures are the first line of such targets [30, 35, 36], while the cerebellum is almost beyond the scope of interest.

Meanwhile, considering the statement on GABA role in AS, the question arises if the cerebellum could be a source of additional strengthening of GABA-ergic mechanisms provoking AS?

To answer such a question means observing the evolution of epileptic activity induced as a result of GABA inhibition disturbance caused by penicillin – antagonist of CABAA receptors under conditions of cerebellar ES. Penicillin – induced activity is recognized as closely resembling SWD, and systemic administration of epileptogen to feline cats is a hallmark of such resemblance [37]. Nevertheless, foci induced via application of relatively low dosage of penicillin solution upon brain cortex start their activity from purely negative spikes and demonstrate step by step development of particular discharge (spike) component – such as primary slow-wave, positive component (PPC) following spike [38]. Such a “minor” characteristic of epileptic discharges might be informative for antiseizure action of antiepileptic drugs [38, 39]. We were interested in two phenomena, which are underlying with GABAergic mechanisms. Namely, GABA-mediated inhibition contributes to the profound postsynaptic inhibitory potential of substantial length in penicillin-induced foci, which underlay slow-wave genesis [20–22, 26, 37]. The genesis of PPC reflects local inhibitory (“surround inhibition”) [40–42] barrier activity, which does not permit to expand epileptic activity along with the neural tissue.

Hence, our investigation aimed to observe the specificity evolution of interictal penicillin-induced cortical focal activity, emphasizing slow-wave characteristics, and spikes PPC under paleocerebellar electrical stimulation (ES).

## 2. MATERIAL AND METHODS

The experiments performed on 15 male cats, weighing 2,5-3,5 kg under acute experimental conditions. Procedures involving animals and their care were conducted according to University guidelines that comply with international laws and policies [European Community Council Directive 86/609, OJ L 358, I, December 12, 1987; National Institute of Health Guide for Care and Use of Laboratory Animals, US National Research Council, 1996].

All animals underwent tracheostomy and skull trepanation performed under ether anesthesia. Points of pressure in a stereotaxic frame and all soft tissue dissection zones were infiltrated with 0,5% of novocaine solution, and this procedure repeated every 2,0 h. Tubocurarine (“Orion,” Finlandia, 0,2 mg/kg, i.v.) was injected, and after that, the cats were artificially ventilated. Stimulative nichrome bipolar electrodes (outer diameter 0,12 mm, interelectrode distance 0,2 mm) were inserted to the paleocerebellum (declive, pyramis) under visual control and fixed to the skull with quick-drying dental cement.

The dura mater was dissected 2,0-2,5 h from the moment of the cessation of ether anesthesia, and filter paper (2,0x 2,0 mm) was soaked with *ex tempore* prepared sodium benzylpenicillin solution (16,000 IU/ml) applied to the posterior sigmoidal gyrus. The indifferent electrode was then placed in nasal bones. Monopolar EEG registration performed using electroencephalograph 4- EEG-3 type (FSU). Electrical stimulations (ES) of the cerebellum were done with an electrostimulator ESU-1 (FSU) (100-300 Hz, 0,25 ms, 150-250 µA, duration of trial-3-7 s). The interval between ES was not less than three minutes. Only histologically controlled locations of electrodes were included in observation. Control cats were false-stimulated [43].

## 3. RESULTS

Fifteen to twenty min after the moment of application of the epileptogen upon the cat’s cortex (posterior sigmoidal gyrus), single spikes with an amplitude of 1,5-2,2 mV and the frequency of generated discharges was 30 to 50 per min (Fig 1, A).

**Fig.1.**
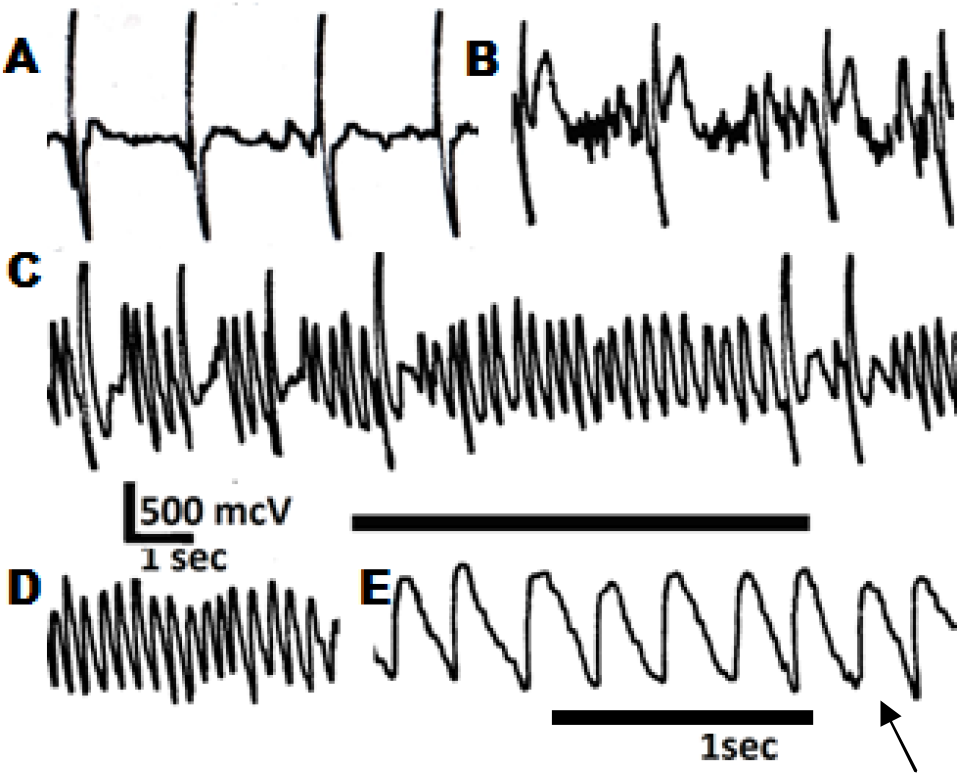
Effects of paleocerebellar ES upon penicillin-induced foci in brain cortex of myorelaxed and artificially ventilated cats. A- 17,5 min after the moment of application of penicillin sodium salt (16,000 IU/ml) to the posterior sigmoid gyrus; B-1,0 min after the cessation of 2-d ES and 5,5 min after A; C-5th ES; D-, E- 1,0 min after the moment of 7th ES; With an arrow, a small amplitude spike preceding a sharp wave marked. Parameters of ES: 100 Hz, 0,25 ms, 250 mcA (period of ES marked with a solid line at C).

Not regular and small PPC was registered at this period in control observations with false stimulations (as an example, look at fragment A, which demonstrates the period before ES start). More regular PPC with amplitude than 25% of the spike’s total magnitude followed with slow wave appeared in the next 15,0 – 20,0 min in the control group. Meanwhile, the development of pronounced (up to 50% of the total magnitude of discharge) PPC of discharges followed with pronounced – up to 0,6 mV slow wave was induced in 1-3 ES (Fig. 1, B). This time, the discharge amplitude diminished to 1,3-1,8 mV, while the frequency of generation of discharges reduced to 15 - 45 per min. During the next 2-4 ESs, the suppression of spikes along with the appearance of slightly distorted sinusoidal waves (3,5-3,75/sec) with an amplitude of 0,45-0,80 mV was registered (Fig 1, C). Consequent 2-4 ES were enough to suppress all spiking activity. This effect was observed in 5 out of 7 experiments, and in all of them, spikes were substituted by relatively regular (3,5-3,75/sec) sinusoidal waveforms with a constant amplitude of 0,6 to 0,9 mV (Fig. 1, D, E). Not regularly, small amplitude spikes preceding wave were identified (look at E, the arrow pointed). It should stress that any SWD appearance was seen in control observation - false-stimulated cats with spontaneous declining focal activity.

New local application of penicillin (16,000 IU/ml) in the zone in which sinusoidal waves were present (Fig. 2, A) caused an initial decrease of their amplitude (Fig. 2, B) with the appearance of spikes in 1,0-3,0 min (Fig. 2, C). In the next 1,0-4,5 min, spikes’ amplitude reached their maximal value (Fig. 2, D). Remarkably the PPC was absent in newly appeared discharges.

**Fig 2.**
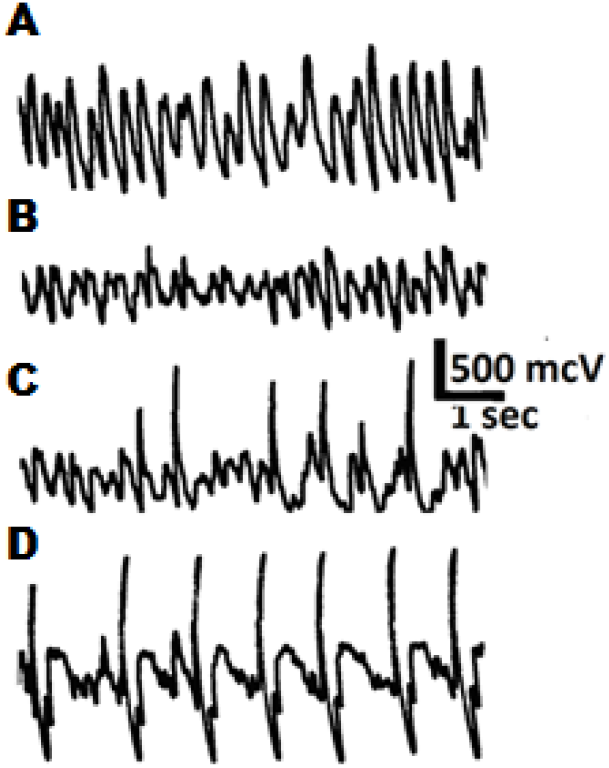
Effect of penicillin application in the zone with sharp spindle-like waves generation. A- 1,5 min from the moment of 8-th ES of paleocerebellum; B- 0,5 min, C- 2,5 min, and D- 5,5 min from the moment of the new application of benzylpenicillin (16,000 IU/min). (Adopted from [43])

## 3. DISCUSSION

Hence, delivered results revealed the precipitation of not-in burst organized prolonged slow-wave activity with the frequency of 3,5-3,75 Hz in the zone of cortical penicillin-induced foci. Such activity is close to such one characteristic for absence SWD and induced via paleocerebellum ES.

As far as cortical administration of GABAA antagonists (penicillin) precipitated seizure activity with SW pattern [44, 45], the spontaneous occurrence of SWD was expected in the course of the natural decline of penicillin-induced foci activity. Such type of evolution did not take place in our observations but provoked with paleocerebellar ES. Obtained results correspond with the data on the precipitation of SWD (3/sec) in the rat brain cortex on the focal penicillin-induced foci model caused by paleocerebellar ES [47]. They correspond also with the theory on the leading cortical role in SWD generation [9, 46] and with the general conception on the role of GABA-ergic mechanisms in SWD precipitation.

Gained data permitted to state the next properties of described regular slow waves:

-waves rows are restricted to the zone of primary penicillin application and did not impact intact cortical zones;
-waves shape is asymmetrical with a sharp increase and slow decline of amplitude;
-suppression of slow waves caused by additional penicillin application – the fact in favor of their genesis via activation of CABA-dependent mechanisms of their appearance. Vice versa, the activation of local inhibition caused by cerebellar ES is also supported by the induction of pronounced PPC as far as its development reflects the state of local surround inhibition [40–42]. Precipitation of pronounced PPC was an immediate result of first-second stimulations.

Hence, it might be supposed that influences coming to the forebrain from stimulated paleocerebellum can precipitate EEG signs of AS in the cortical GABA-A mechanism zone disturbed with local penicillin application. This effect of ES is promoted by the increase of GABA-ergic inhibition in target neuronal chains as far as investigated components of seizure discharged (slow wave, PPC) reflect the heightened state of GABA inhibition and developed in parallel with the suppression of spike discharges [43, 47, 48].

The involvement of GABA strengthened inhibition in observed effects of transformation of epileptic activity is supported with clinical magnetic resonance spectroscopy data revealing the heightening of GABA-ergic mechanisms in the brain caused by cerebellar transcranial magnetic stimulation (TMS) [49, 50]. The authors established that cerebellar TMS followed by an average increase of EEG synchronization in the theta–bandwidth, accompanied by the rise in regional GABA level in the prefrontal cortex [50].

Existing literature data coming from different methodical approaches to the clarifying cerebellar role in AS are in line with the gained data:

### 3.1. Tonic onic and phasic effects of cerebellar ES

The prolonged poststimulative character of SWD provocation is in favor of the tonic nature of AS facilitation, which might realize via extrasynaptic δ-containing GABAA receptors [51] located both in the thalamus [52] and in the brain cortex [53, 54]. It should stress that extrasynaptic tonic inhibition is necessary to induce AS. Thus, knockdown of eGABAAR by genetic methods prevented precipitation of AS both in the GHB model and spontaneous AS in GAERs rats [19]. GABA elaborated in cerebrospinal fluid in the course of high-frequency (200 Hz) ES of the cerebellum [55, 56] might also contribute to eGABAAR activation.

Another mechanism on such stimulation realized via the strengthening of GABA synthesis from glutamate elaborated from glutamatergic cerebellar efferents and accumulated in thalamic targets in the course of serial cerebellar ES has been assumed by Gornati SV and Hoebeek FE [see 57].

Opposite to tonic, phasic influences upon the CTC network are realized transsynaptically [58]. Hence, the disruption of the CTC network via prolonged depolarization of thalamic neurons and dysfacilitation of epileptogenesis, as a result, is recognized as the most probable antiseizure role played by cerebellar nuclei in SWD suppression [59]. Pharmacological (gabazine) local stimulation of cerebellar nuclei neuronal activity followed with SWD suppression favors the proposed conception [60].

Different effects of phasic and tonic effects of cerebellar ES upon SWD are in correspondence with suppressive action of intrathalamic administration of glutamate receptors agonists upon SWD, while similar administration of GABA-ergic compounds (gamma-vinil-GABA, tiagabin) stimulate SWD generation in rats with genetic forms of AS [61, 62].

### 3.2. Imaging of absence seizure network

It is challenging to apply positron emission tomography (PET) to investigate ictal phenomena and their interpretation [63]. That is why the case of absence status epilepticus is of particular interest in clarifying its pathogenesis with [18F]FDG-PET method [64]. The authors concluded that thalamus activation, hypometabolism in frontal, parietal, and posterior cingulate cortices, and hypermetabolism in the cerebellum favor the maintenance of absence status (3-4 Hz rhythmic delta waves in EEG). Earlier, [65] delivered similar data based on [18F]FDG-PET in a patient with absence status. Namely, the authors pointed to bilateral thalamic hypermetabolism and frontal cortex hypometabolism. BOLD data on GSWD activity in patients with juvenile myoclonic epilepsy revealed the net cerebellum involvement with the negative relationships between the thalamus, cerebellum, frontal and sensorimotor-related areas [66–68].

Using EEG/fMRI, the source of generalized SWDs determined in patients who suffered from resistant idiopathic generalized epilepsies [69]. The authors analyzed 36 AS and revealed that peak blood-oxygen-level-dependent (BOLD) response was reached maximally in six seconds from the AS EEG onset with a simultaneous time-schedule for the temporal lobe. The stable peak of BOLD response in the cerebellum registered one sec after the thalamus’s peak favors that ictal AS induce disturbances encompassing the cerebellum. Following the classical statement, seizures arise from the cerebral cortex, spreading to other structures, including the cerebellum [70]. Nevertheless, cerebellar structures’ primary role in seizures generation is also suspected [71] and disclosed on pentylenetetrazol-modeled seizures in Zebrafish [72].

### 3.3. Impulse activity of cerebellar cells in seizure suppression

PCs are activated during the occurrence of SWD, as well. High voltage spindles, recorded epidurally from the rat sensorimotor neocortex, correlated with single or multiple unit activity in the cerebellar cortex and deep cerebellar nuclei [73]. Later on, [74] reported that slow synchronized oscillations in the neocortex drive slow oscillations in the cerebellar cortex. Specific changes of extracellular spike trains of cerebellar nuclei of Cacnala tg mice, another genetic absence model, precipitated in the course of SWD appearance, favor cerebellum involvement [75].

Up to 26% PC demonstrated an increase of complex spike activity and rhythmicity during generalized SWDs in homozygous tottering mice [76]. Those effects are better pronounced in the cerebellar cortex’s lateral parts, which receive inferior olive inputs. Interestingly, the bilateral inferior olive lesion was produced by systemic administration of the neurotoxin 3acetylpyridine, followed by a proconvulsant state-specific for strychnine-induced seizures myoclonus [77].

New data coming from the optogenetic experimental approach must favor increased impulse activity. Either PC [78] or nuclei cells [60, 79] resulted in the suppression manifestations of temporal lobe epilepsy induced with intrahippocampal kainic acid administration.

That is why cerebellar cells’ increased impulse activity induced in SWD development assumes that backward cerebellar efferent influences aim to seize and suppress seizure discharges. However, in the case of absence electrogenesis, such a situation is different from the temporal epilepsy model, and backward cerebellar influences could maintain absences manifestations.

## 4. CONCLUSIONS AND PERSPECTIVES

Hence, the presented data favors the modulative and triggering role of the cerebellum in AS electrographic manifestations. The cerebellum’s pro-absence seizure potential may at least partially contribute to cerebellar DBS’s inconsistency in experimental conditions and patients with resistant epilepsy [36, 71, 80–82].

Some perspectives comprise parallelism between minor AS behavioral manifestations and functionality of the cerebellum extended beyond motor control. Thus, the lately established cerebellar role in cognition [83–85] corresponds to the well-known disruption of consciousness and disturbing informational processes in WAG/rij rats [5, 6, 86, 87].

AS prevalently occurred at an early age (5 - 12 years), and manifestations lessened with the aging. A similar schedule of cerebellar neurons degeneration was observed in aging [88]. Such parallelism, together with discussed data on the functional state of cerebellum and precipitation AS manifestations, permits the assumption that cerebellar influences severely modulate the absence-prone CTC network. Such a belief is in line with the data [89], who described a default mode network in patients with AS, which extended beyond the CTC network and included striatum and reticular pons structures.

## Abbreviations

AS: absence seizures
BOLD: blood-oxygen-level-dependent (response)
CAE: childhood absence epilepsy
CTC: cortico-thalamo-cortical (network)
DBS: deep brain stimulation
EEG: electroencephalogram
eGABAAR: extrasynaptic gamma aminobutyric acid type A receptors
ES: electrical stimulation
GHB: γ-hydroxybutyric acid (butyrolactone)
fMRI: functional magnetic resonance imaging
GABA: γ-aminobutyric acid
PC: Purkinje cells
PPC: primary positive component
RT: reticular (thalamic) nucleus
SWD: spike-wave discharges
TMS: transcranial magnetic stimulation.
[18F]FDG-PET: fludeoxyglucose (18F) – positron emission tomography.

## ACKNOWLEDGMENTS

The authors express their thankfulness to Professor Gilles van Luijtelaar and Professor Coenen A.M.L. for their valuable critical remarks and recommendations.

## Authors contribution

LS created the conception and wrote the manuscript. POleh and POlesya analyzed the data, contributed to final design and edited the manuscript.

## Ethics approval

The work was approved by the Ethics Committee of Odesa National Medical University and conforms to the European Communities Council Directive 24 November 1986 (86/609/EEC; National Institute of Health Guide for Care and Use of Laboratory Animals, US National Research Council, 1996).

